# New water and air pollution sensors added to the Sonic Kayak citizen science system for low cost environmental mapping

**DOI:** 10.1101/2020.11.03.366229

**Authors:** Amber GF Griffiths, Joanne K Garrett, James P Duffy, Kaffe Matthews, Federico G Visi, Claire Eatock, Mike Robinson, David J Griffiths

## Abstract

Sonic Kayaks are low-cost open-source systems for gathering fine-scale environmental data. The system is designed to be simple to fit on to standard kayaks or canoes, and full instructions have been made available for anyone wishing to build their own. The first version included temperature sensors and a hydrophone for recording underwater sound. Here we outline the design and integration of two new sensors, for underwater turbidity and above water air particulate pollution. All sensors record continually, and the GPS location, time and date are also stored for every data point, allowing fine-scale environmental data mapping. The data being collected by the sensors is sonified (turned into sound) in real-time, allowing the paddler to hear the data as they are collecting it, making it possible to locate and follow interesting occurrences. We present proof-of principle data for the first time for all the sensors, demonstrating how the system can be used for environmental mapping, and discuss potential future applications and adaptations. We believe the Sonic Kayak system offers particular promise for citizen science and environmental activism, as well as allowing professional researchers to gather data that was previously difficult or impossible to obtain.

## Introduction

The Sonic Kayak is an open source technology project for gathering the environmental data for making fine-scale maps of marine, estuarine, river or lake environments. The equipment is designed to fit onto any model of kayak. The Sonic Kayak Version 1 (Griffiths et al. 2017) was equipped with temperature sensors and a hydrophone, recording the water temperature every second and underwater sound continuously, together with the GPS coordinates, time and date. The data is also sonified in real-time through an on-board speaker, allowing the paddler to hear the data as it comes in and to follow interesting occurrences, for example temperature gradients (sonification the process of conveying information by using non-speech sounds; Dubus & Bresin 2013, Hermann et al. 2011).

Since publishing the original Sonic Kayak design we have had requests to add a turbidity sensor to measure the cloudiness of water caused by particulates. The requests were from people wanting to monitor algal blooms in an EcoPort, monitor cyanobacteria for a water company, and take water quality readings for seaweed farming. Similar applications could include mapping farm run-off or sewage outflows. We were also interested in measuring particulate pollution in the air low over the water, caused by boat engines and industry. This type of air pollution is not well studied, but has the potential to affect water users like swimmers or kayakers as well as wildlife such as waterbirds or air-breathing cetaceans (e.g. Rawson et al. 1995).

Low cost open source and open hardware environmental sensors are proliferating and becoming more reliable, for example the water quality sensor by Bas Wijnen et al (2014) has been technically verified against professional equipment, and the Luftdaten air quality project is now producing data globally from open source particulate matter equipment (https://sensor.community/en/). The Sonic Kayak system sets itself apart from these projects as the environmental data is geolocated and can be collected while moving, allowing very quick mapping without the need for multiple sets of equipment. A similar initiative, the Smartfin (https://smartfin.org/) is close in its aims, but is specific to surfboards and stand-up-paddleboards (as the electronics are embedded within a fin), currently only measures temperature, and is not open source so is not available for people to build or develop themselves (Brewin et al. 2020).

This article outlines the design of the new turbidity and air quality sensors and how to fit them into the Sonic Kayak, together with the new data sonification approach for these sensors. In addition we present proof-of principle data collected using Sonic Kayaks, including maps for water temperature, underwater sound, water turbidity and air particulate pollution. Fine-scale mapping of environmental data in difficult-to-reach areas like estuaries and close to coasts is an exciting step for professional researchers, however we believe the system also offers particularly interesting opportunities for citizen science and community-driven environmental activism. The Sonic Kayak system is highly flexible, and could equally be used on land, for example to collect particulate pollution data while walking or cycling around a city.

### Hardware and Software

#### General design considerations

Version 1 of the Sonic Kayak (Griffiths et al. 2017) was designed with the electronics, sensors, and two speakers located at the front of the kayak. Since then, we worked with an accessible kayak club (Access Lizard Adventure https://www.lizardadventure.co.uk/category/access-lizard-adventure/) to redesign the Sonic Kayak system for people with visual impairments, to enable greater independence on the water. As part of that redesign (Version 2), we moved the electronics and sensors to the back of the kayak, and switched from cabled speakers to a single waterproof bluetooth speaker that is located in the middle of the kayak where there is usually a recess for holding a water bottle. These changes were largely to reduce the risk of entanglement in the cabling, and of course, this is beneficial to every paddler. The new design also had the benefit of being quicker and easier to attach to the kayak as the main electronics box now sits in the storage compartment at the back of the kayak and simply clips onto any available d-ring or strap that the kayak model has. During the Version 2 redesign we also improved the waterproofing using a watertight electronics box (IP66/68 rated, meaning it is resistant to full submersion of up to 1.5m of water for up to 30 minutes) fitted with marine-grade cable glands.

The addition of the new turbidity and air quality sensors for Version 3 have brought their own challenges. Two of the key design considerations for the new water turbidity sensor were (i) to keep it at a fixed depth to ensure consistent data, and (ii) to reduce the light from above water that could affect the data collected. To achieve this we built a structure that attaches to the side of the kayak, allowing the paddler to easily lift it out of the water when needed. With previous versions we were aware that the temperature sensor also suffered from not being at a fixed depth; in this version we improved this issue by fixing it to the same rig as the turbidity sensor. The air quality sensor, by definition, needed to be open to the air, which makes full waterproofing impossible. Because of this it needed to be as low cost as possible, and separate from the main electronics, meaning that if the kayak capsized this sensor could be destroyed without affecting the rest of the equipment. Together this means that the Sonic Kayak version 3 system is mainly housed in a waterproof box on the back of the kayak, with a small external box containing the air quality sensor on top, a hydrophone trailing from the back of the kayak, and a rig strapped using velcro to one side handle of the kayak holding the temperature and turbidity sensors, together with a separate waterproof bluetooth speaker that can be placed anywhere on the kayak (Figures 1 and 2).

**Figure 1.**
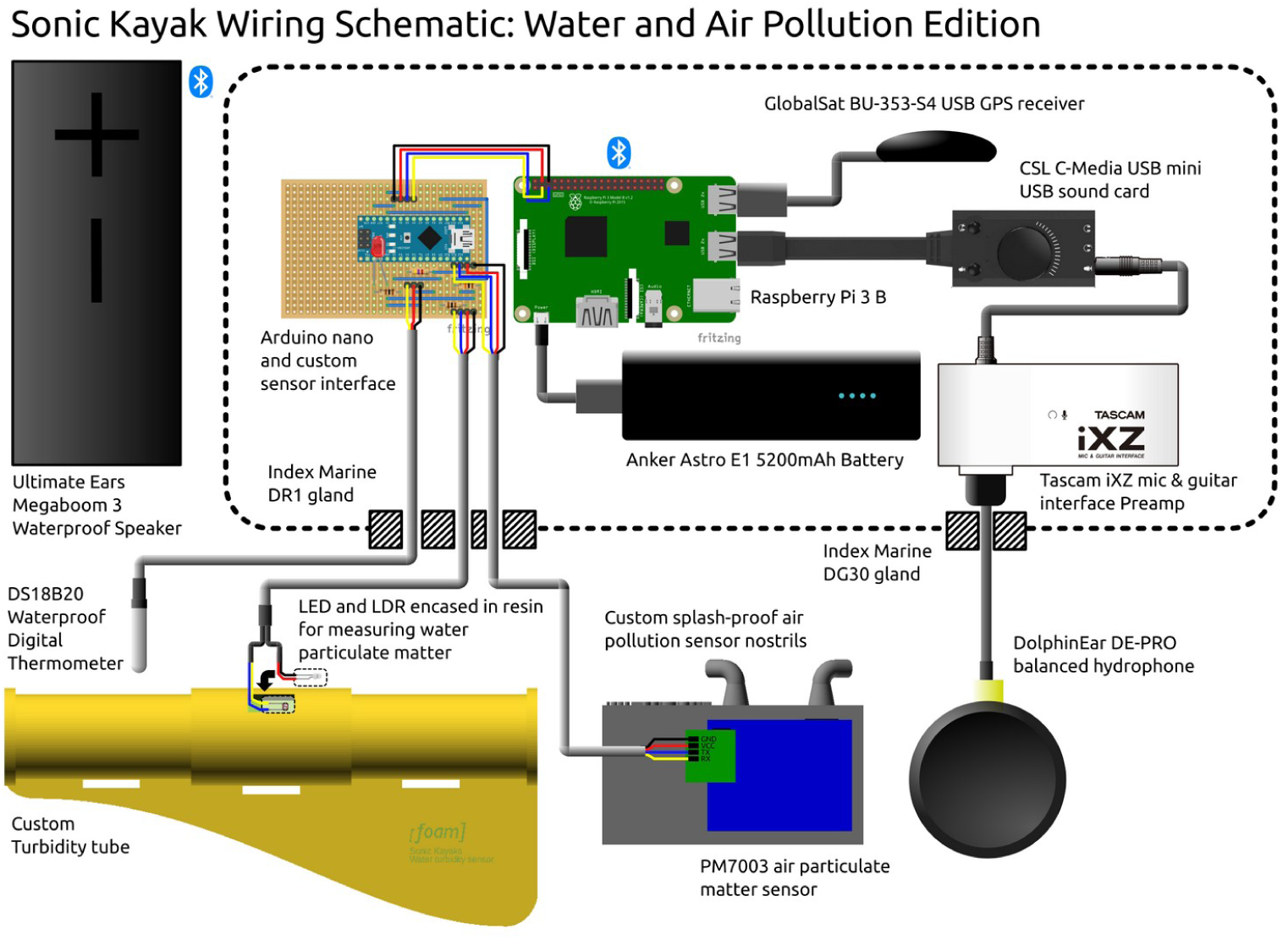
Schematic for a Sonic Kayak system, showing the different components including sensors, and how they fit together.

**Figure 2.**
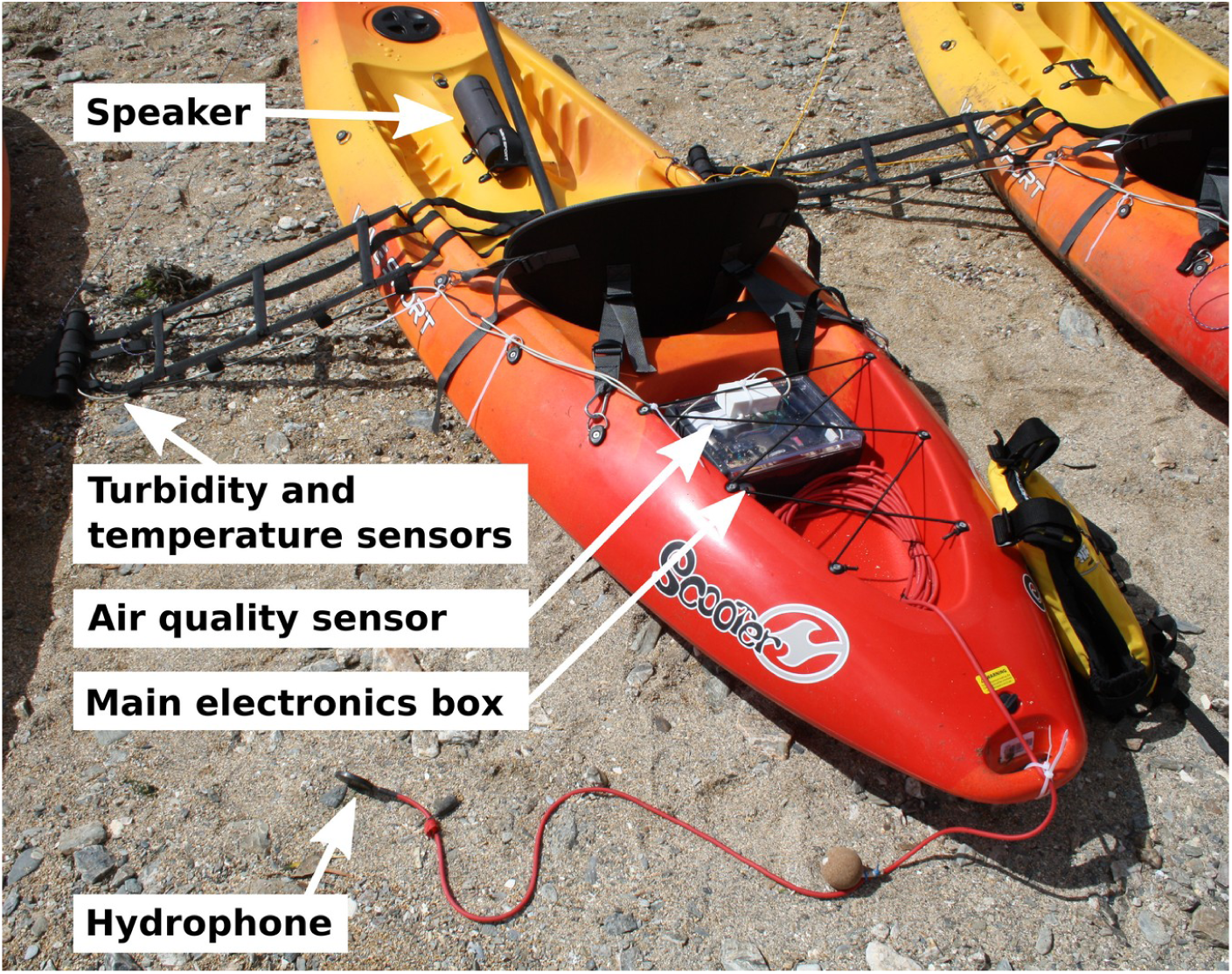
The Sonic Kayak version 3.

Detailed plans for building a complete Sonic Kayak (version 3) can be found in the project wiki https://github.com/fo-am/sonic-kayaks/wiki, and the new sensor design decisions are described in more detail below. The software is open source and available on GitHub https://github.com/fo-am/sonic-kayaks.

#### Particulate sensor design

Low cost sensors for measuring air quality (specifically particulate matter) have been implemented very successfully for citizen science in many settings, and most comprehensively by Luftdaten (https://sensor.community/en/). Reliable linear relationships have been demonstrated for various low-cost particulate matter sensors tested against professional research grade equipment (e.g. the TEOM™ Continuous Ambient Particulate Monitor from Thermo Scientific), however the low-cost sensors do appear to routinely over-estimate particulate levels (Badura et al. 2018). Low cost PM2.5 sensor measurements can be affected by humidity, mainly from fog as opposed to rain, as the water particles are similar in size to the pollution particles (Jayaratne et al. 2018) -which is of course a concern for use on water.

Of the four low-cost models of particulate sensors tested by Badura et al. (2018), PMS7003 (manufactured by Plantower) and SDS011 (manufactured by Nova Fitness, and used by the Luftdaten project mentioned above) appear to perform best in terms of data replicability and comparisons with research-grade equipment, however humidity levels were shown to affect SDS011 such that the data was less comparable with the research-grade TEOM data the higher the humidity. This did not appear to be an issue with PMS7003. Temporal drift can also be a problem with low-cost PM sensors either from dust accumulation or the degradation of electrical components, but Bulot et al. (2019) saw no evidence for drift over time for the PMS7003 sensor. Because of these issues, we decided to work with the PMS7003 sensor instead of the more widely used SDS011 sensor.

We designed a custom 3D printable housing for the PMS7003 sensor to protect it from rain and splashes (Figure 3), including downward pointing tubes aligned with the inflow and outflow of the sensor which allow air in/out, but will stop ingress from water (unless submerged). Files for 3D printing are available via GitHub https://github.com/fo-am/sonic-kayaks/tree/master/hardware/3dp (pm-box is as pictured in Figure 3, pm-box-home-print is a version with wider air input/output nostrils that is also better suited to printing on a standard extrusion printer).

**Figure 3.**
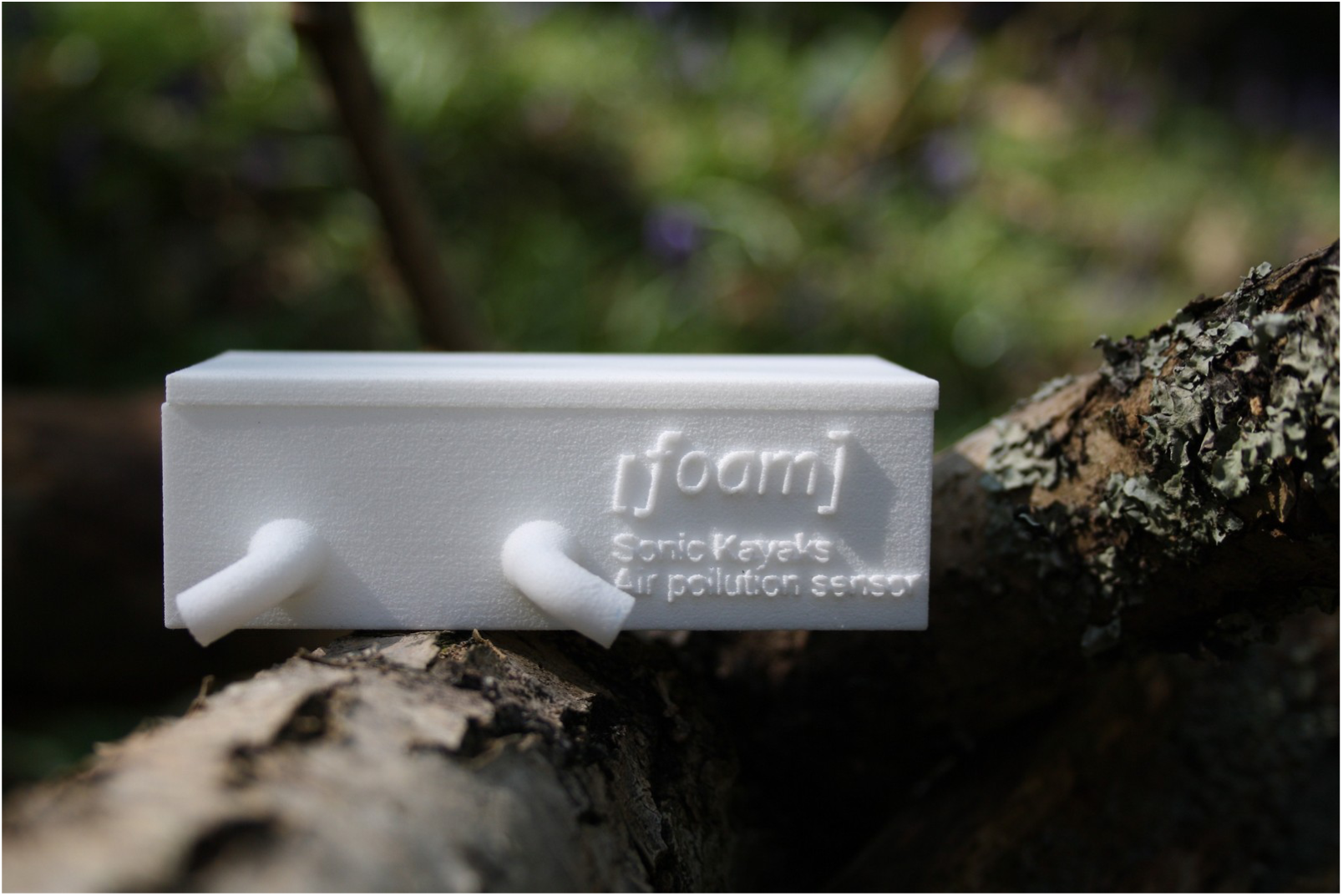
Custom housing for PMS7003 air quality sensor.

The PMS7003 sensor outputs particulate matter concentrations for PM1/2.5/10 at ‘standard particle’ (μg/m3; the concentration corrected to a standard atmospheric pressure) and ‘under atmospheric environment’ (μg/m3; the concentration in the air as it is when the sample was taken, which can be useful for example when taking measurements at high altitudes as air pressure changes the concentrations of gasses), and the number of particles of each size in 0.1 litres of air. For the purposes of this paper we have used the standard particle levels, having checked that they do not differ substantially from the atmospheric equivalent values at sea level.

We believe the best approach is to consider only relative values rather than to expect accurate absolute values, to primarily use the system for studying spatially localised particulate matter concentrations, and where possible to take multiple readings for the same area to confirm replicability. If absolute particulate matter concentrations are needed, we recommend calibrating the air quality sensor using professional-grade equipment that includes a drying mechanism for the air sample, or against an Automatic Urban and Rural Network (AURN) station (UK) or equivalent (unfortunately there are no AURN stations in the >100km of land west of Plymouth in the UK, which is the region that our trials took place).

#### Turbidity sensor design

Turbidity sensors give a measurement of the amount of suspended solids in water – the more suspended solids, the higher the turbidity (cloudiness) of the water. The most basic approach to measuring water turbidity is to use a Secchi disk. These are plain white or black and white circular disks that are lowered into the water, and the depth at which the disk is no longer visible is an approximate measure of the cloudiness of the water. This is a great low-key approach, but the result is affected by other factors such as the amount of daylight (Preisendorfer 1986). More accurate equipment tends to use an enclosed light source and a light receptor, with the water placed in between, so that the amount of light that reaches the receptor from the light source gives a reading of how turbid the water is. There are several pre-existing publications on how to make open source turbidity sensors (e.g. Román-Herrera et al. 2016 and the Public Lab turbidity sensor prototype https://publiclab.org/wiki/turbidity_sensing). For the Sonic Kayaks, we sonify sensor data in realtime and record the data continuously. This means we need to make a sensor that logs real-time data and provides a digital output that allows it to be integrated into the existing Sonic Kayak kit, as opposed to a system where you take a one-off sample of water and run it through a separate piece of equipment in a laboratory. We based our design on Bachler’s (2019) prototype, which uses a Light Emitting Diode (LED) and Light Dependent Resistor (LDR) housed within a piece of hose pipe - the more cloudy the water is, the less light from the LED will reach the LDR.

We designed a custom 3D printable turbidity tube, with a reinforced nylon webbing rig that attaches to the kayak handle using velcro. This means that the sensor is kept at a reasonably fixed depth (approximately 60cm below the waterline), but can easily be lifted out of the water where it is too shallow, and is very quick to attach/remove. The rig also provided an opportunity to mount the temperature sensor at the same fixed depth. The turbidity tube itself was printed using selective laser sintering for strength, with a matte black nylon plastic to block light, and was designed with slatted end caps to allow water to flow freely through while keeping debris out of the tube, as well as to further minimise light entering the tube. The files for 3D printing are available via GitHub https://github.com/fo-am/sonic-kayaks/tree/master/hardware/3dp (turbid-tube and turbid-cap and suitable for laser sintering prints, or turbid-tube-home-print and turbid-tube-home-print are better suited for low cost extrusion printing). The end caps are held in place with silicone sealant to make them easy to remove for cleaning. The LED and LDR were set into small clear resin blocks for waterproofing (Figure 4a), orientated over component-sized holes drilled into the turbidity tube, and fixed in place with matte black Sugru to block out the light (Figure 4b). The cables run up the nylon webbing rig such that there is no pressure on any of the joins (Figure 4c).

**Figure 4.**
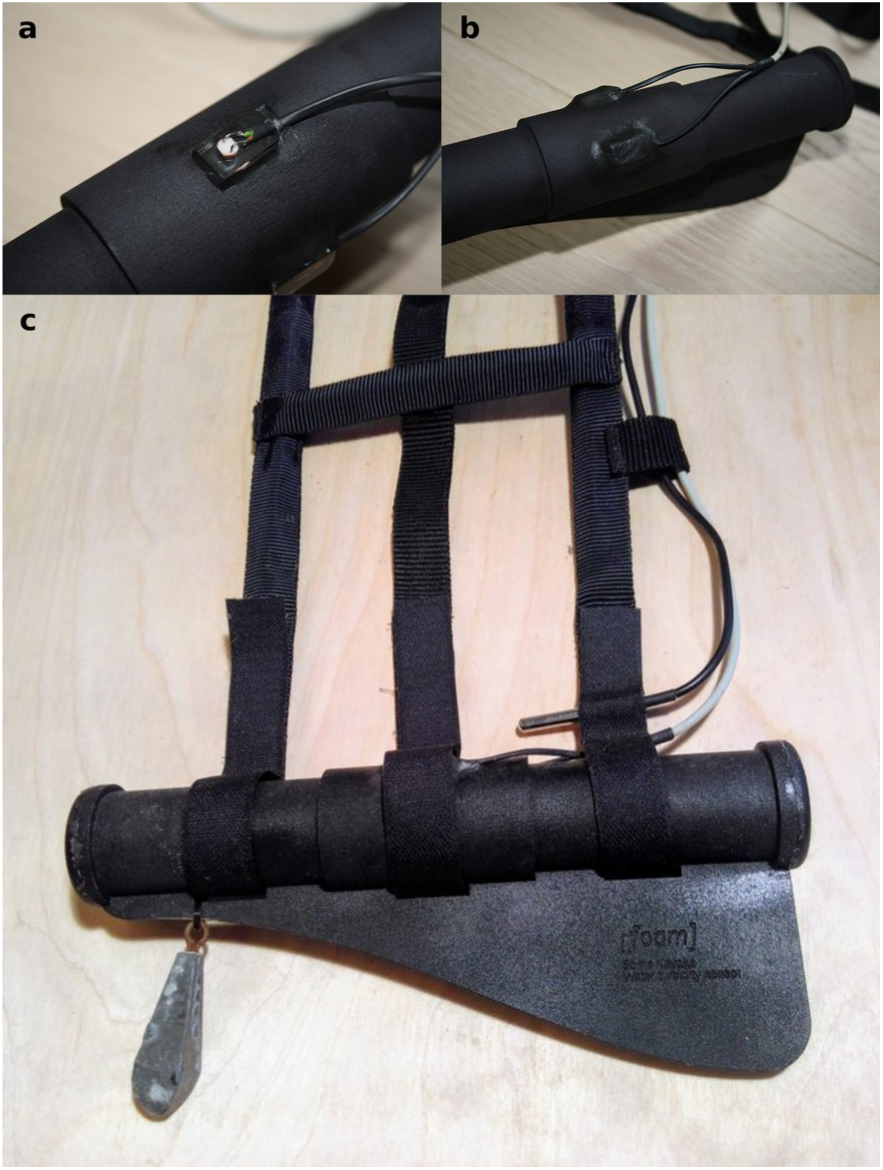
(a) Electrical component (LDR) set in resin block attached to the turbidity tube, (b) Both electrical components (LED and LDR) attached to the turbidity tube and coated to block light, (c) The completed turbidity sensor attached to the nylon rigging with velcro straps, with the temperature sensor just above, and an optional weight.

The data output from the turbidity sensor is not completely straightforward. LDRs decrease resistance with light intensity so when more light hits the sensor, the less resistance there is and the higher the voltage reading is, resulting in a higher numerical output. The numerical output is related to the voltage coming in, with an analogue to digital conversion (10 bit) applied such that 0V = 0 and 3.3V = 1023. If required, it is possible to do a lookup from the specific LDR sensor curve data to work out the voltage from the numerical output. As with the air quality sensor, the turbidity sensor can easily be calibrated using professional research equipment, and without calibration it is suitable for obtaining relative values and identifying spatial areas with high/low turbidity.

The light levels above water may influence the readings despite our efforts to block extraneous light, however tests where the kayak was paddled under the shade of low bridges indicated that this level of change in above water lighting conditions was not noticeable on the output, and similarly two different Sonic Kayak systems paddled around the same area on different days and at different times of day under different weather conditions produce remarkably similar results (Figure 5).

**Figure 5.**
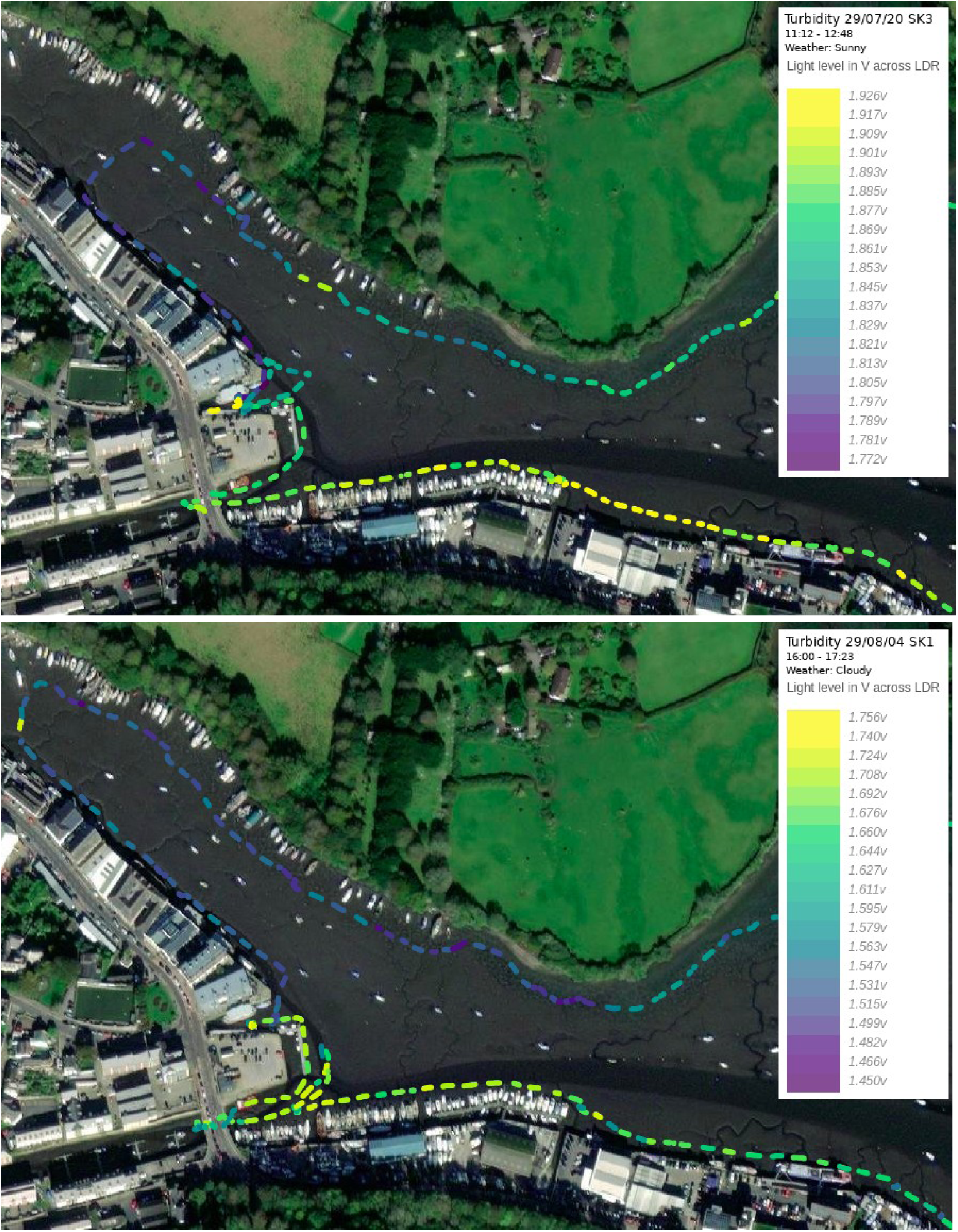
Comparison between two turbidity sensor recordings on different days with different weather conditions at high tide. Absolute differences in voltages are due to slight differences in hardware construction and resistor tolerances. Samples recorded in clearer river water allow more LED light to pass through than in the muddy estuarine areas. Passing underneath the bridge in the lower left does not cause significant changes in light sensor readings on either trip.

### Sensor Integration

#### Software

The particulate, turbidity and thermometer sensors were connected to an Arduino Nano microcontroller, attached to the main Raspberry Pi system. The reasons for using an Arduino Nano in addition to the Raspberry Pi were to (i) provide analogue to digital conversion for the turbidity sensor which the Pi lacks and (ii) to free up a connection (UART) on the Pi (this connection is required for the PMS7003 and the GPS). The Arduino connects to the various digital and analogue interfaces for each sensor, collects the data and provides it to the Raspberry Pi in a convenient form.

The 10 bit analogue to digital converter (ADC) on the Arduino uses the battery power level as its reference voltage, and so is susceptible to fluctuations on the power supply coming from other parts of the circuit. We found that the PMS7003 causes high frequency interference (either from the motor air pump or laser) which affected our turbidity measurements. We were unable to alleviate this interference with either the addition of extra bypass capacitors or via software filtering. In order to sidestep this issue we found we could use the command interface on the PMS7003 to put the sensor to sleep for ten seconds while reading the turbidity sensor. A small delay is required to allow the PMS7003 to fully power down before reading a clean value from the ADC, and ten seconds is an adequate length of time to allow air to be drawn into the particulate sensor to allow accurate periodic readings.

Once read by the Raspberry Pi over an i2c serial interface, the data is (i) appended to a log file stored on a USB stick in CSV format to be analysed later and (ii) sent to the sonification system, alongside error conditions so the paddler is aware of changes in the data or problems with any of the sensors.

Python scripts read and log sensor data and send it to Pure Data for sonification. The GPS is interpreted by an audio zone map to indicate areas for sampling (see below) by playing indicator sounds when you cross into polygons drawn on a map before setting off. Due to the unpredictable nature of Bluetooth radio transmission, a watchdog process detects problems that can affect the hydrophone recording (e.g. speaker out of range) and can restart and reconnect to the speaker. Underwater recordings are split into 26Mb (5 minute) mono WAV files and written to USB stick including sequence and trip IDs embedded in the filenames.

All software is available via GitHub (https://github.com/fo-am/sonic-kayaks), and Figure 6 shows a schematic of how the various software components interact.

**Figure 6.**
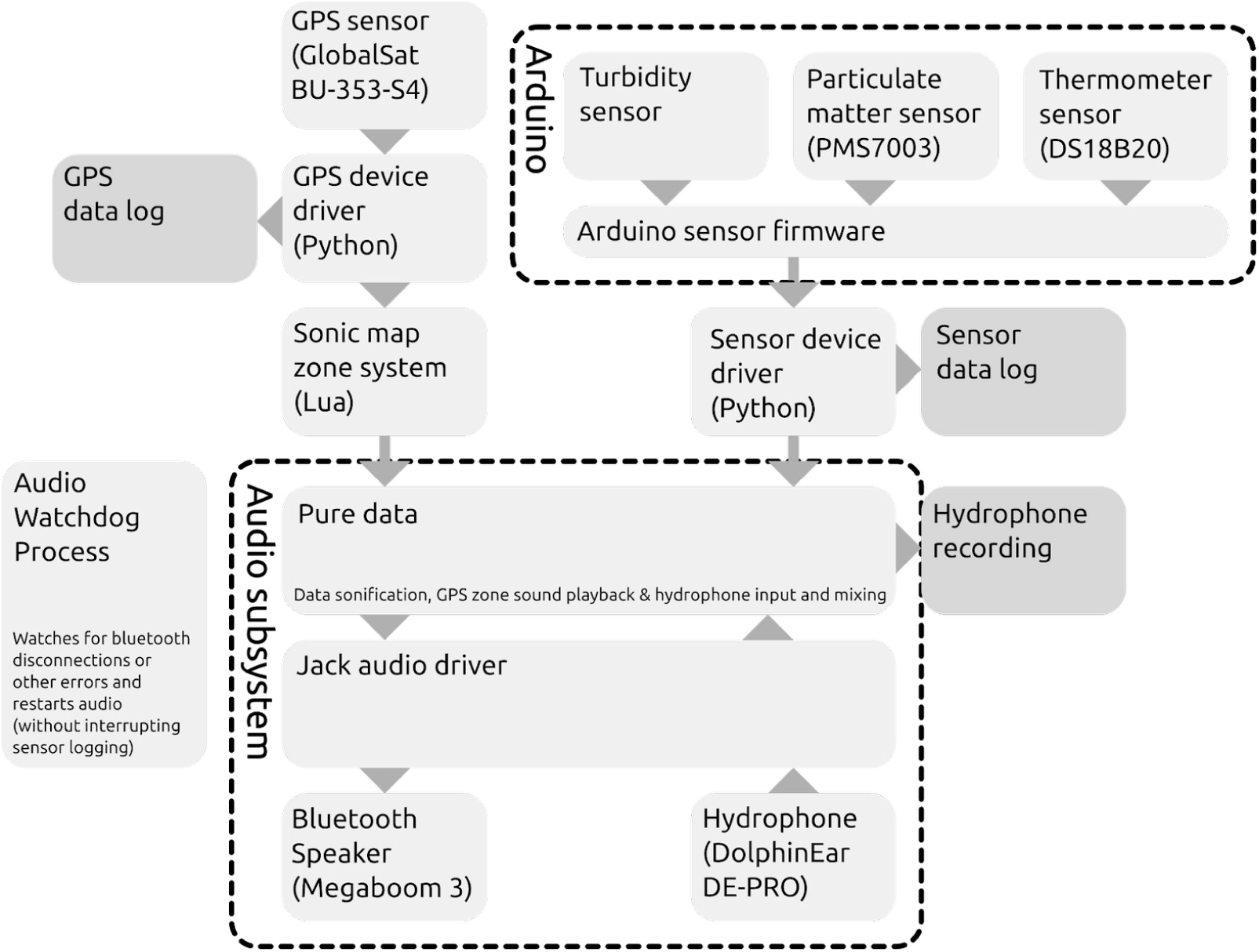
The Sonic Kayak software setup for sonification and storage of sensor data.

### Data sonification

The Sonic Kayak version 1 sonification consisted of a live-feed from the hydrophone so the paddler could hear the underwater sounds, together with a simple synthesised tone for the temperature sensor which went up in pitch in higher temperature water, and down in pitch in lower temperature water (only making a sound when the temperature changed, not continuously). Adding two new sensors might seem straightforward, but in terms of the sonification, this required three separate sounds (representing temperature, turbidity, and air quality) that could be distinguished easily from each other, sounded good together and with the hydrophone feed, and didn’t overwhelm or annoy the listener/paddler. To handle the task of sonifying multiple data streams in different ways more easily, we built a custom modular sonification system in Pure Data that allowed us to use different synthesis and sample manipulation techniques on separate sounds.

To guide our decision making, we made four separate types of sonification for the set of sensors, and produced an online survey to see which people preferred. We also made mock environmental data sets where the data for each sensor was either rising/falling/rising then falling/falling then rising, and tested survey respondents to see whether they could tell what was happening with the environmental data just from the sonification. The questions and survey response data is available in full on Zenodo, from 49 survey participants (https://zenodo.org/record/3923743#.X0j2OHWYVH4). We chose simple synthesised sounds which scored second for favourability (mean score = 3.41, on scale of 1-5), and scored first for interpretability (73% correct answers). A demo track with these sonifications is available here: https://archive.org/details/skdemo_20200915.

Another important aspect of the audio feedback for citizen science uses is error detection. As trips can last multiple hours, the paddler needs to be aware of problems as soon as they happen - rather than returning to base to discover some or all of the data is unusable. If a sensor stops working the system uses synthesised speech to warn the paddler immediately that there is a problem - this is much easier to keep track of than lights or a visual display would be in the water. For some sensors, there are specific error states that need to be reported - such as GPS “no fix”, where not enough satellites are acquired the system makes an audible “ping” warning that the positioning is currently bad quality.

### Proof-of-Principle Data

#### Data collection

To demonstrate the potential of the Sonic Kayaks, we performed tests in two locations in Falmouth Bay, Cornwall, UK (Figure 7), which is a Special Area of Conservation (SAC) and a Special Protection Area (SPA; for birds). The Bay is used by cargo shipping, cruise ships, fishing and recreational boating. The first location was Falmouth Harbour (29th July 2020). To act as a good case study we picked a highly variable area/route(s) - this test covered an estuary from the river through to more open sea, including places where people live on houseboats, areas under agricultural fields, an industrial dock, town, and marinas of various sizes. For this test we took three systems out, each with turbidity, temperature, and air quality sensors. The second test location was the Helford River (7th August 2020), which is a flooded river valley that is a Site of Special Scientific Interest and within the Cornwall Area of Outstanding Natural Beauty. This area is typically thought of as very quiet, clean and beautiful, attracting holiday makers and leisure craft users (particularly yachts and kayaks). For this test we took one system out, with turbidity, temperature, air quality and hydrophone sensors. The locations included the habitats eelgrass and Maerl beds which are features protected within the SAC designation (JNCC 2015).

**Figure 7.**
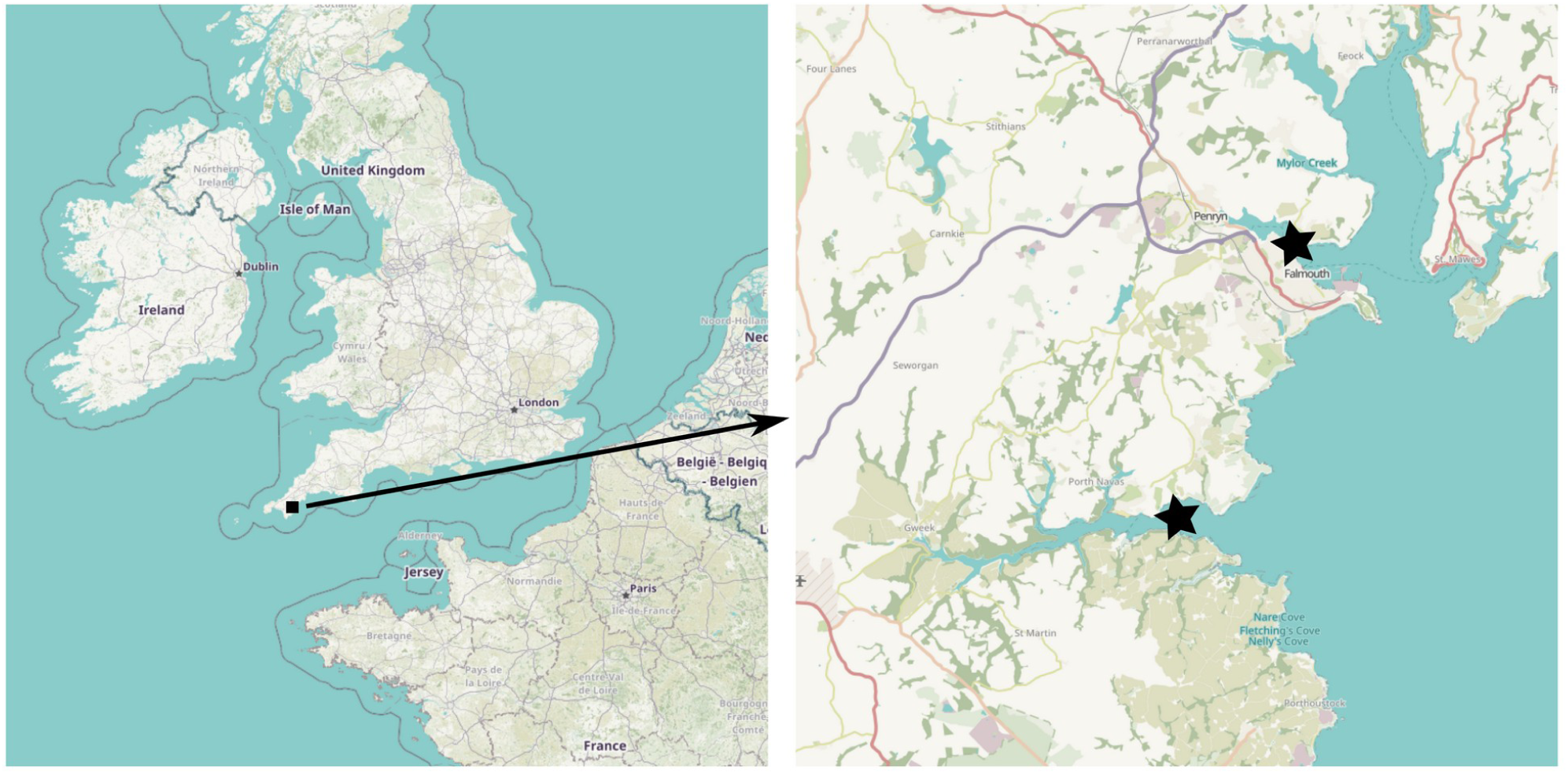
Map of two test locations, Falmouth Harbour to the North and Helford River to the South (base map from Open Street Maps).

#### Data analysis

Data from the temperature, turbidity, and air quality sensors do not need any post-processing, however the underwater sound data from the hydrophone is more complex.

The hydrophone data is collected continuously, but to obtain good quality recordings ideally it is best to avoid any noise, including the noise from paddling, or drips coming from the paddles. As such, we decided to stop kayaking at semi-regular intervals, roughly every 200m, to gather a clean sound sample. To facilitate this, we made use of a feature built into the original Sonic Kayak system that allows you to take a map of an area and define spatial zones, then attribute sounds to those zones that will automatically play through the speakers when the kayak enters the zone (triggered by GPS). We placed zones approximately 200m apart around the path that we were planning to paddle the kayaks, so that an alert ping sounded to prompt us to stop for a minute. This was developed from the previous approach which was more *ad-hoc* and where samples were not taken periodically (Griffiths *et al*., 2017). This method allows for a more systematic approach to sampling.

#### Sound analysis

The .wav files were viewed (in spectrogram mode) in the freely available open-source sound software Audacity (version 2.2.2) to identify aurally periods of time without paddling noise. The start and end time of these sampling periods were recorded (at 1 second time resolution). These samples were analysed in RStudio (version 1.2.5001; RStudio, Inc.) with R version 3.6.3 (R Development Core Team, 2020) with custom scripts adapted from Merchant *et al*., (2015) and using the package “tuneR” (version 1.3.3; Ligges 2018). All code for sound analysis is available via GitHub (https://github.com/fo-am/sonic-kayaks/tree/master/analysis).

A correction factor was calculated by recording sine waves (at 10 Hz, 100 Hz, 1kHz and 10 kHz) of known voltage (measured using a Amprobe PM51A Pocket Digital Multimeter) through the entire system (Tascam iXZ pre-amp, TechRise USB sound card and Raspberry Pi) and comparing the output with the known input. This was then interpolated to provide a correction factor per 1 Hz. The hydrophone sensitivity, provided by the manufacturer as a range (−180 to −183 dB; although not calibrated), was specified in our processing as −181.5 dBV μPa-1. The correction factor and hydrophone sensitivity allows absolute sound values to be calculated. However, these should be interpreted with some caution as the hydrophone sensitivity is not calibrated.

To explore how the sound level varies with frequency (and to apply the frequency dependent corrections), fast Fourier transforms were applied to the sound samples in 1 s segments (Hann window, 50% overlap). Several metrics were calculated. The overall sound level for the full frequency range effectively recorded with the hydrophone was represented by the broadband sound pressure level (SPLRMS). This was calculated per 1 s by summing the square pressure values within the range 20 Hz - 19 kHz.

Variation in sound levels with frequency (pitch) are represented by third octave levels (TOL). An octave represents a doubling in acoustic frequency from the lower limit to the upper limit of the band (frequency range) and third octave bands are 1/3 of an octave wide. The third octave bands are described by the centre frequency and the bandwidth increases with increasing frequency. Third octave levels were calculated by summing the square pressure values within the range of third octave bands with centre frequencies from 25 Hz - 16 kHz.

Sound levels were also calculated for wider frequency ranges intended to approximately represent a range of boat sizes. Large commercial ships are represented by the lowest frequency range of 20 Hz - 100 Hz. For example, a cargo ship of 173 m length produced the loudest sounds in the third octave bands of 25 Hz or 50 Hz, depending on speed (Arveson & Vendittis 2000). Large boats, such as fishing boats and ferries, are represented by the range 101 Hz - 1 kHz. The loudest frequencies from a fishing vessel recorded in China, were generally in this frequency range (Zilong *et al*., 2018). Further, the whale-watching vessels recorded in Cholewiak *et al*., (2018) were loudest in the third octave band of 200 Hz, while fishing vessels were loudest in the third octave bands with centre frequencies 200 - 325 Hz. Small boats, comprising mainly recreational vessels, are represented by the frequency range 1.001 kHz - 10 kHz. Recordings of small boats have found peaks in this range (Jensen *et al*., 2009) and at ∼1.6 kHz and higher (as well as loud sound levels at low frequencies) (Sarà et al. 2007). Square pressure values within these ranges were summed together. The square pressure values were passed to the mapping script, averaged over space (time may vary) and then converted to decibels as the final step.

Below we present the broadband sound level; large ships, big boats and small boats frequency ranges; and the 125 Hz third octave level. This 125 Hz band is, along with the 63 Hz band, an indicator of shipping noise in the marine strategy framework directive (MSFD; Tasker et al., 2010). Previous research found the 125 Hz band to be louder in Falmouth Bay than the 63 Hz band (Garrett et al., 2016).

The noise floor of the system was estimated by leaving the Sonic Kayak system to record for five minutes in a quiet space out of the water. The analysed recordings were above the noise floor for >99.9% of the time for all presented measures, with the exception of the 125 Hz third octave level (∼99.7%).

#### Data visualisation

Data collected with the different sensors on the Sonic Kayak are time-stamped and therefore matching them up is easily achievable. This means that each data point has a position in space as provided by the on-board Global Navigation Satellite System (GNSS). The following processing and figure creation was conducted in the R software environment (R Development Core Team 2020), with the following packages: cowplot (Wilke 2019), dplyr (Wickham et al. 2020), ggplot2 (Wickham 2016), lubridate (Grolemune & Wickham 2011), purrr (Henry & Wickham 2019), raster (Hijmans 2020), readr (Wickham et al. 2018), RStoolbox (Leutner at al. 2019), sf (Pebesma 2018) & tidyr (Wickham & Henry 2019).

Regular 50m wide grids were created spanning the extent of GNSS data for the Falmouth Harbour and Helford sites. These hexagon polygons were then clipped to a mean high water line freely available from the Ordnance Survey (https://osdatahub.os.uk/downloads/open/BoundaryLine). Next, the sensor data were joined (using time as a common variable) to the GNSS data to create a spatial point dataset. These points were then intersected with the regular hexagon polygons, and the mean values of the sensor measurements were calculated and assigned to the hexagon. This process was repeated for each of the sensors. Not all of the sensors were simultaneously recording throughout the whole kayak trip (e.g. in the case of turbidity and air quality, the sensors alternated). Absent data were treated as NA values and had no influence on the calculated means. If a hexagon intersected points only with NA values, then it was assigned an NA. These hexagons appear hollow in Figures 8-11, indicating the path of the kayak, but the absence of data. Lastly, spatial data was projected to British National Grid (EPSG 27700) and overlaid on mosaicked aerial images (Sources: Esri, DigitalGlobe, GeoEye, i-cubed, USDA FSA, USGS, AEX, Getmapping, Aerogrid, IGN, IGP, swisstopo, and the GIS User Community). All data used in the visualisations are available via Zenodo (https://zenodo.org/record/4041588) and the code used is available via GitHub (https://github.com/fo-am/sonic-kayaks/tree/master/mapping).

**Figure 8.**
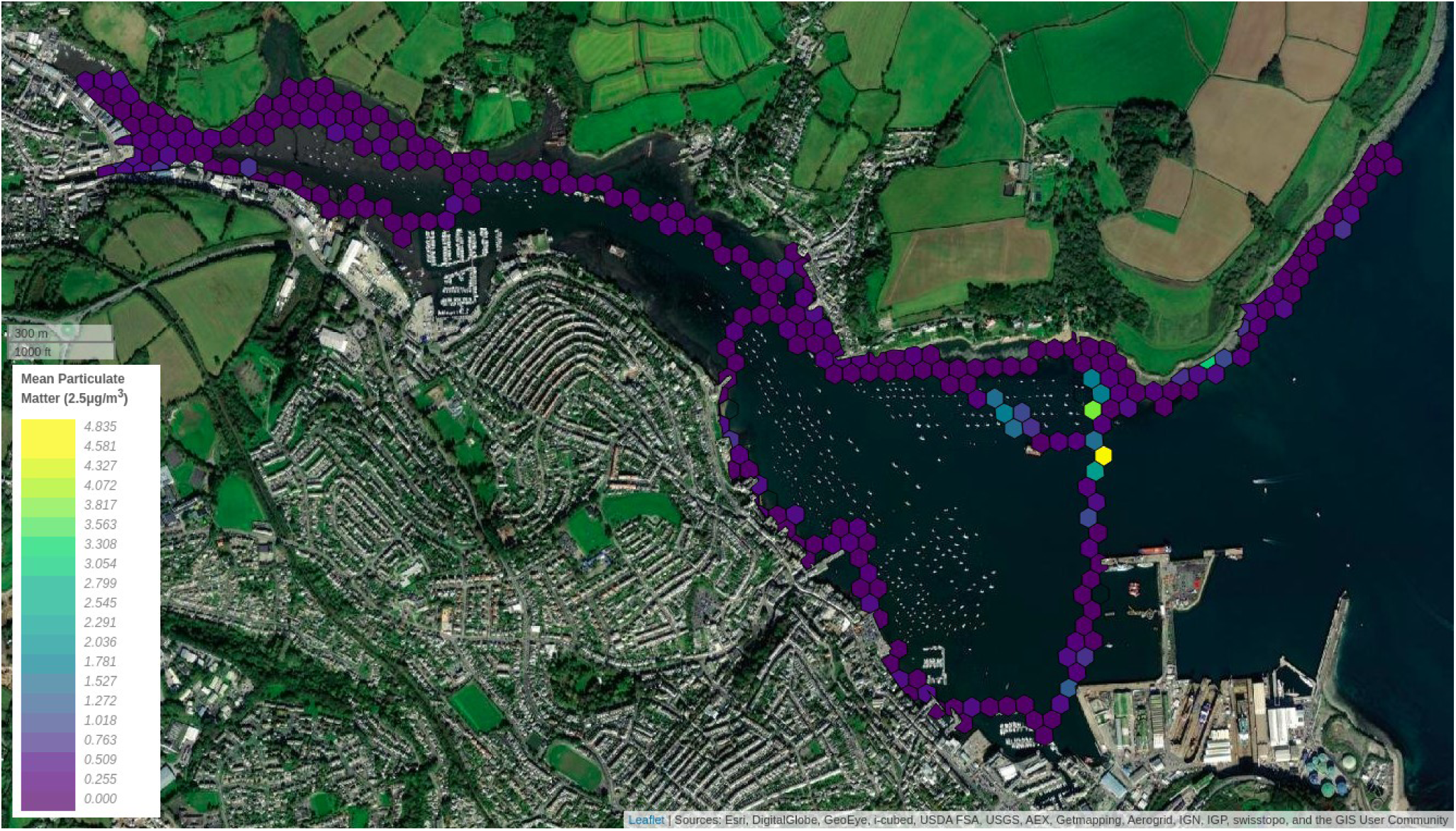
Falmouth Harbour particulate matter data (PM2.5).

#### Data interpretation

The data visualisations from the test trips give an indication of the potential of the Sonic Kayak system for fine-scale mapping, particularly close to the coast, but also in rivers and lakes.

Spatial and temporal variation are evident, for example the Falmouth Harbour particulate matter map shows a spike in particulate pollution just to the North of where The World cruise ship was moored and running its engines despite being stationary, with the wind coming from the South at the time of sampling (see Figure 8 for PM2.5, and Supplementary Material 1 for PM1 and PM10). This example shows the potential for communities to use the system to highlight poor practice and lobby for change.

The turbidity sensor, which is perhaps the most likely to be problematic as it is not an off-the-shelf sensor, shows good replicability from multiple trips at Falmouth Harbour - with higher levels of water turbidity detected in muddy tidal areas, and lower at a river outflow (Figure 5 for detailed version, Figure 9 for broader geographical area). Water turbidity can be caused by many things, including pollutants like algal blooms or sewage outflows. This system would allow the mapping of the spatial range of such issues which could be of value for environmental protection.

**Figure 9.**
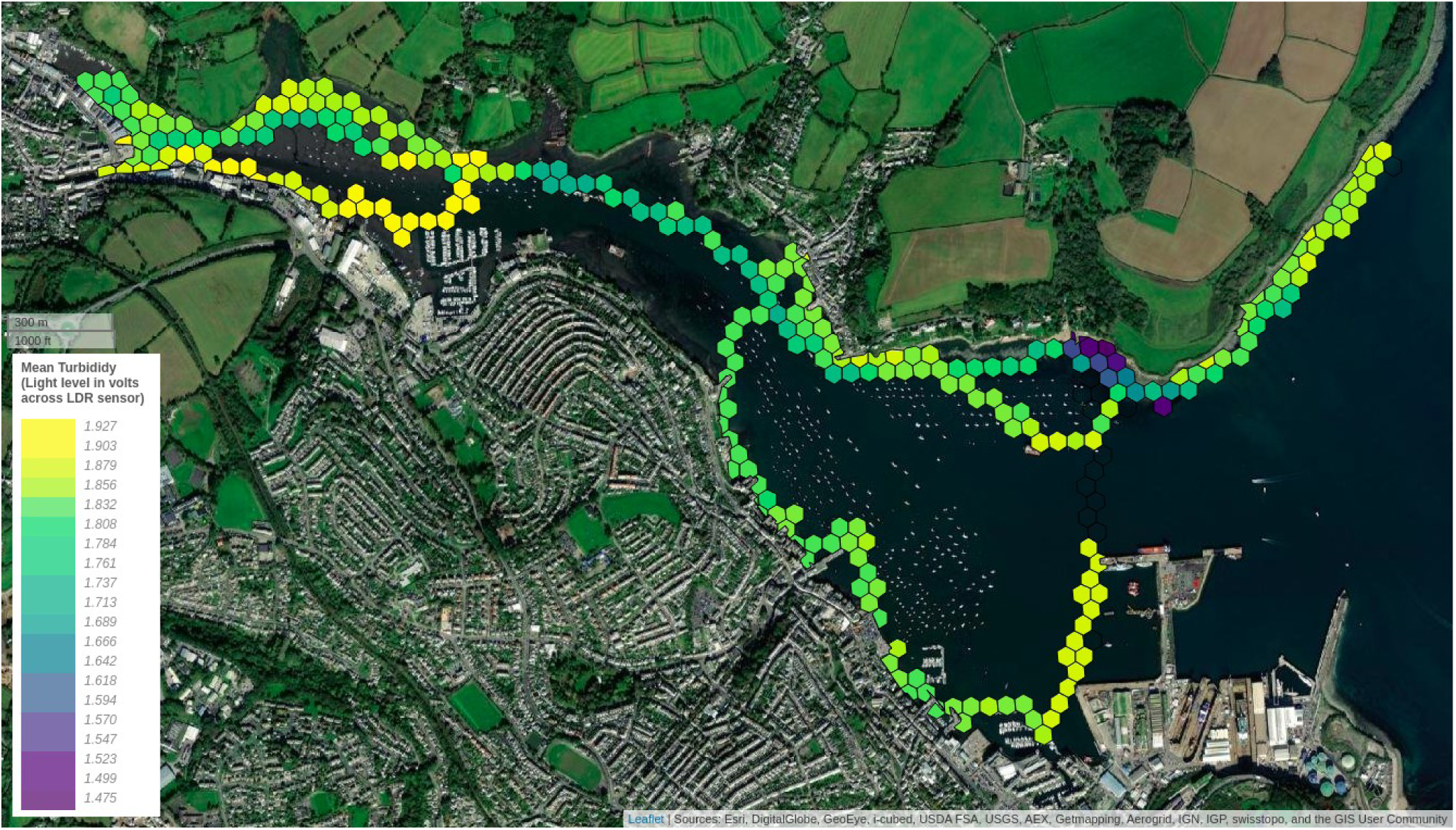
Falmouth Harbour water turbidity.

Sea temperatures are typically measured using static sensors attached to buoys or other structures, or using low resolution satellite observations (Griffiths et al. 2017). Our test data from Helford and Falmouth Harbour show water temperature variation at very fine scales, with warmer water in shallow and sheltered areas and colder water in deeper and more exposed areas (Figure 10 and Supplementary Material 1). Temporal replication of trips could be used to see how water temperatures vary over the year, or even to detect longer term trends caused by climate change.

**Figure 10.**
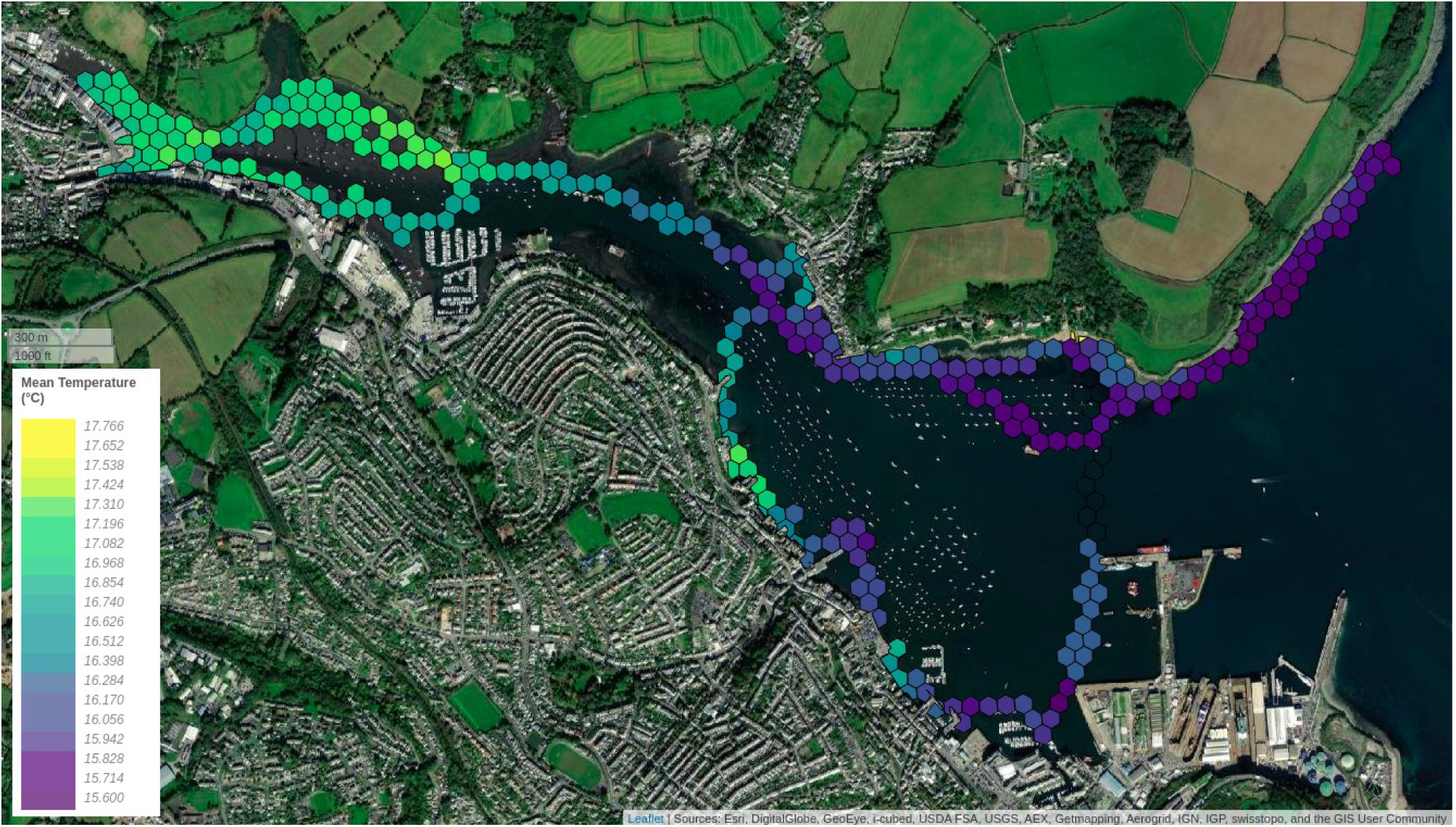
Falmouth Harbour water temperature.

Fine-scale mapping of underwater sound is particularly novel. Underwater sound is usually collected using sensors attached to the seabed, drifting devices, or towed sensors on motored research vessels (Griffiths et al. 2017). The seas around Cornwall have been modelled to have high levels of underwater noise (Farcas et al., 2020). However, the water depth in our Helford test area was no more than 10 m, and such shallow waters prevent the long-distance propagation of low frequency noise such as that generated by commercial shipping traffic. Our results highlight the contribution of higher frequencies to the overall sound levels and the importance of small, recreational vessels in shallow coastal waters (Figure 11). These frequencies are much higher than those typically the focus of international concern and management (Tasker et al., 2010; Farcas et al., 2020; Merchant et al., 2016). The importance of recreational vessels which don’t carry the Automatic Identification System (AIS; the ship tracking system required on larger or passenger vessels), was also highlighted in shallow coastal waters in Denmark (Hermannsen *et al*., 2019). Our example data in the Helford gives an indication of the precise geographical areas that are most affected (towards the South, where the majority of the boats travel, Figure 11 and Supplementary material 1). This type of information could be of value for conservation groups and managers, in deciding whether it is necessary to limit the areas that boats can use, the number/type of boats or speed allowed in a particular area.

**Figure 11.**
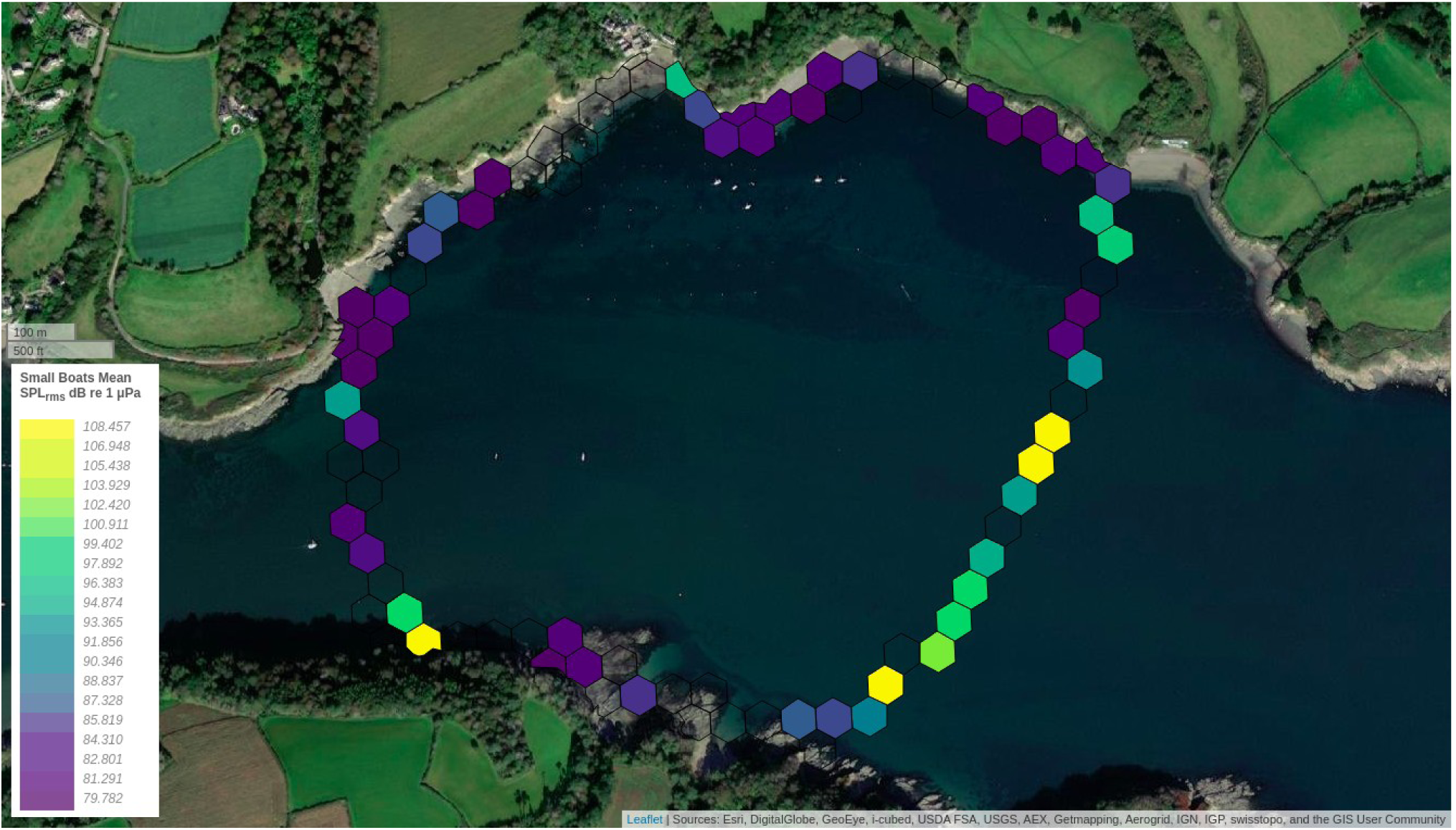
Helford River underwater noise within a frequency range considered indicative of small boats (1 - 10 kHz).

The Sonic Kayak system makes it possible to be very adaptive in how samples are taken. The sonifications allow paddlers to follow interesting data, for example you might hear a temperature gradient and choose to paddle around an area to gather more detailed information in that particular location, or you might hear underwater noise and decide to stop paddling and take a longer sample in that place. It would also be straightforward to modify the sonification approach to notify the paddler when particular thresholds have been crossed, for example ‘safe’ particulate matter concentrations (10 μg/m3 annual mean, 25 μg/m3 24-hour mean are considered important thresholds for PM2.5; World Health Organisation 2005). The kit is very quick and uncomplicated to deploy, meaning that repeat sampling is very feasible. It is as easy to imagine groups of citizen scientists or environmental campaigners gathering data for their own purposes, as it is to see the benefits for professional researchers being able to gather types of data that were otherwise difficult or impossible to obtain (and at a fraction of the cost of typical research equipment). The cost of the whole system, with all sensors (temperature, turbidity, air quality and hydrophone) works out at approximately £1,100 (assuming access to a 3D printer, electronics tools, and other basic workshop tools, as would typically be available in any community Makerspace). Much of this cost is taken up by a hydrophone costing ∼£300 (DolphinEar pro), and the waterproof bluetooth speaker costing ∼ £140 (Ultimate Ears Megaboom 3), each of which may be over or under the required specifications depending on the user’s aims. A full breakdown of indicative costs is given on the project wiki https://github.com/fo-am/sonic-kayaks/wiki.

### Limitations

We recommend that the sensor systems should be used for approximate relative data mapping, rather than for gathering absolute values that would be necessary for certain regulatory or research applications. For professional researchers or those seeking data to guide policy, the system can be of particular use for gathering data to inform the design of more precise secondary studies, or for obtaining more spatially or temporally detailed data following sensor calibration.

The data obtained using the system is very straightforward to handle, with the exception of the underwater sound data which requires more expertise to handle, and requires the manual selection of periods that are clear of paddling noise. It would be possible to modify the system to include on-board data processing for the sound data, to help alleviate this barrier.

In terms of the hardware we are confident that it is robust and highly waterproof, however entanglement and drag remain an issue with the turbidity/temperature sensor rig. It is easy to deal with any entanglement issues while on the water as the rig can be reached from the kayak seat, however if retaining the underwater depth of these sensors is not an issue for the user then the sensors could be fitted close to the hull of the kayak.

### Future potential

We made a video about the project (https://www.youtube.com/watch?v=puLXKj1AVAk) and launched a feedback survey (responses available in full here: https://zenodo.org/record/4032599) to get a broader range of thoughts on the project. Respondents highlighted the diverse potential of the equipment, including for informing decision making in conservation areas, pollution mapping around the coastal sewer outlets and rivers for water companies, land owners and Government bodies, connecting people to the quality of ‘their’ environment and promoting a feeling of engagement/ownership, educational uses with children and others, and monitoring the use and noise distribution of Acoustic Deterrent Devices in open cage salmon farms. One respondent stated that its main use was as ‘a more democratic and participative approach to looking after our natural environments’.

Respondents also suggested future developments could include additional sensors for pH, oxygen levels, and salinity, and some were interested in adding an underwater camera, specifically in one case to capture footage of pelagic marine species and pollution sources (discarded fish nets, waste, etc.) for further qualitative data collection to complement the quantitative data collection. From our perspective we would like to further develop the system to make an off-the-shelf unit that could be purchased at low cost for those who are unable to make their own, and we would like to develop an online mapping portal for hosting, visualising, and sharing the data collected by any users (potentially in real-time).

## Supplementary Material

Supplementary File 1: Data visualisations from test trips.

## Data Availability

Sonification survey data: https://zenodo.org/record/3923743#.X0j2OHWYVH4

Sensor data from test trips: https://zenodo.org/record/4041588

Feedback survey data: https://zenodo.org/record/4032599

Hardware plans: https://github.com/fo-am/sonic-kayaks/wiki

Software: https://github.com/fo-am/sonic-kayaks

Data visualisation code: https://github.com/fo-am/sonic-kayaks/tree/master/mapping

## Acknowledgements

We would like to thank the 49 anonymous sonification survey participants who acted as citizen scientists helping us to decide on the best sonification approach, as well as the 26 participants who gave anonymous feedback on the project as a whole via an additional online survey, some of which has been directly used in the future potential section (consent for use in publication was required as part of the survey).

