## Supplementary file 1 for "New water and air pollution sensors added to the Sonic Kayak citizen science system for low cost environmental mapping"

### Supplementary Material 1: Data visualisations from test trips

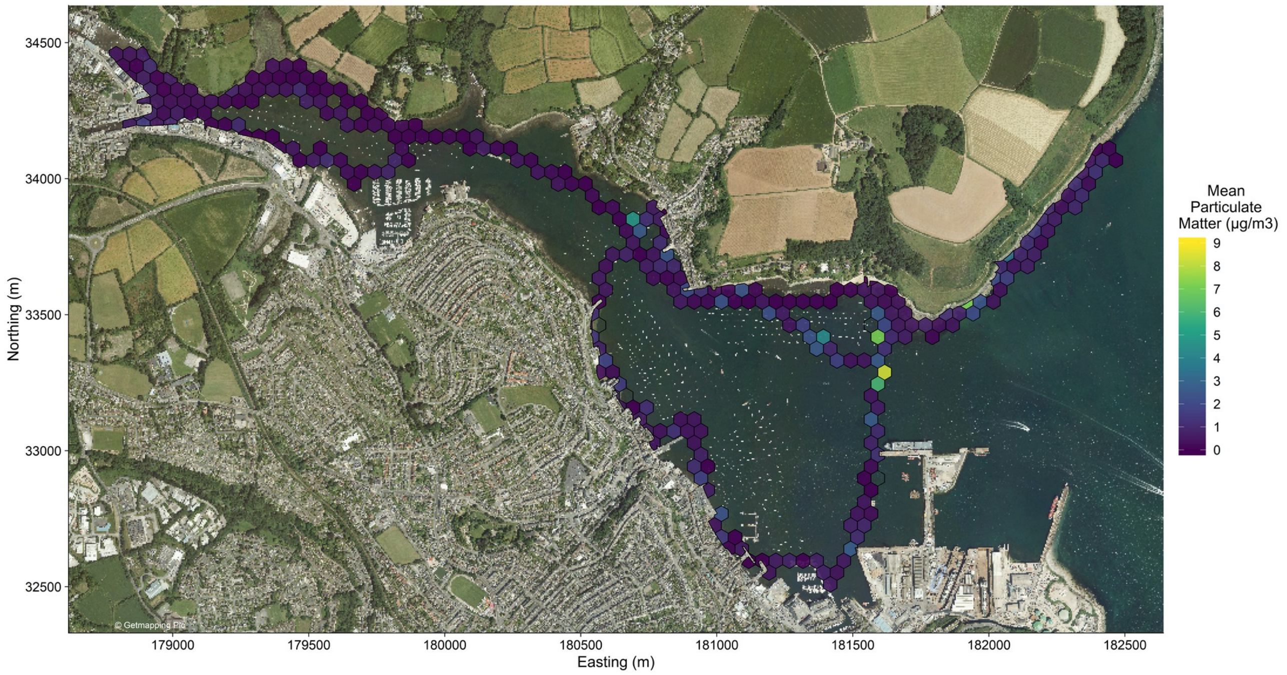


Figure S1: Falmouth Harbour, particulate matter $PM_{10}$

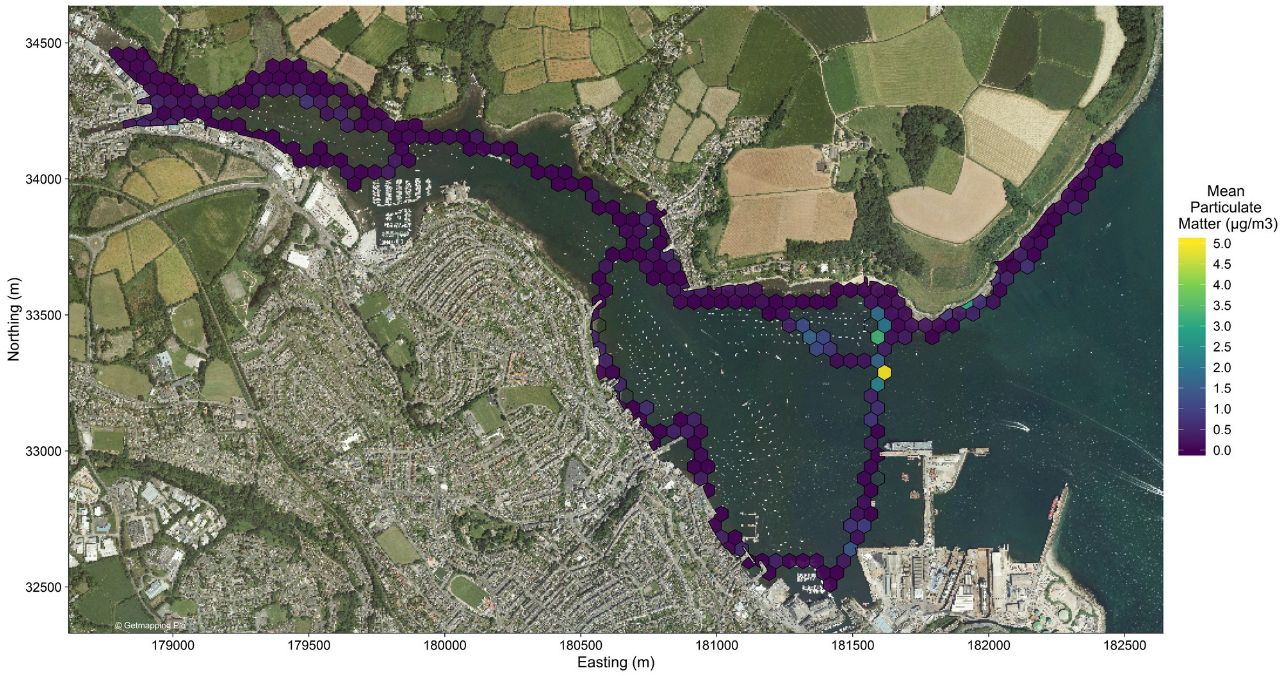


Figure S2: Falmouth Harbour, particulate matter $PM_1$

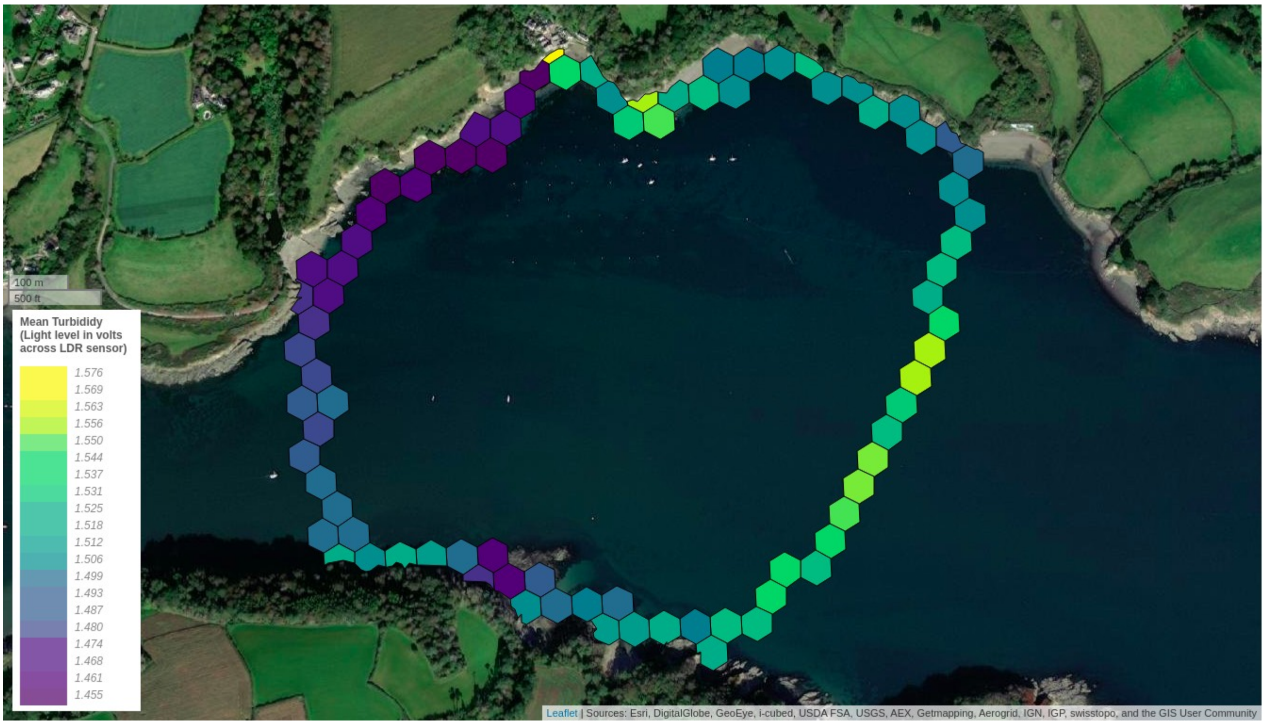


Fig. S3: Helford River, water turbidity.

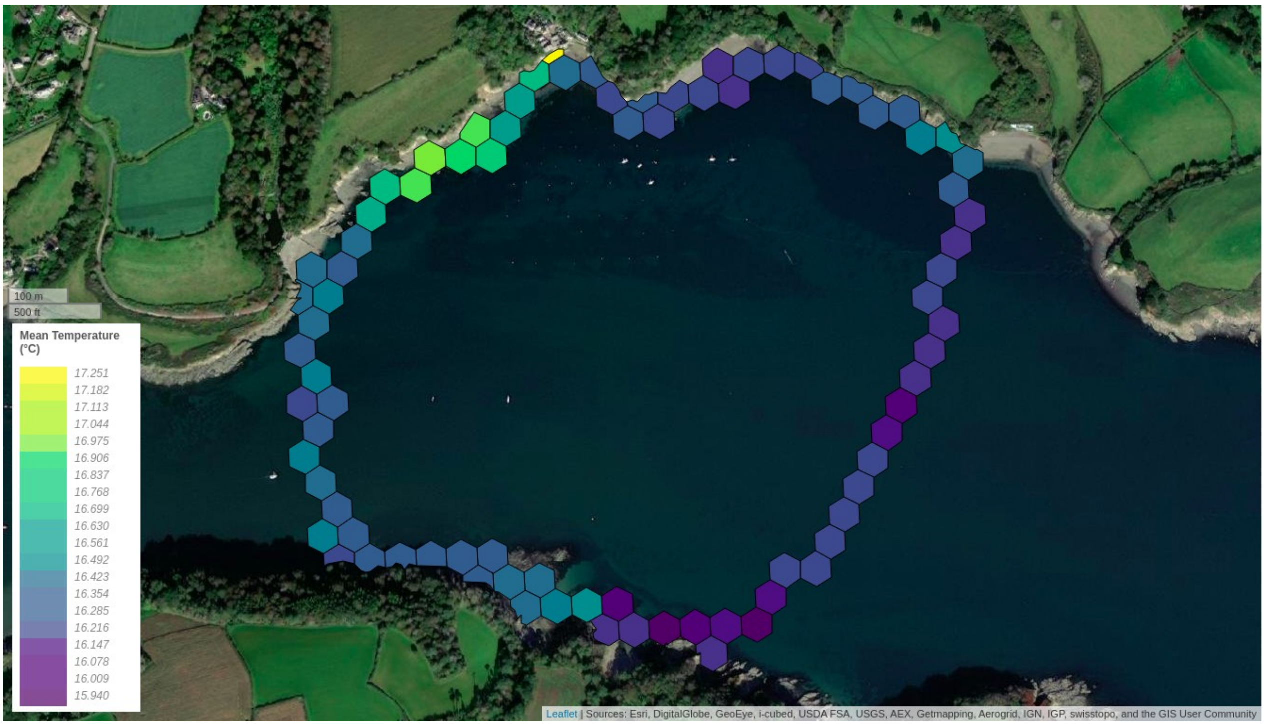


Fig. S4: Helford River, water temperature

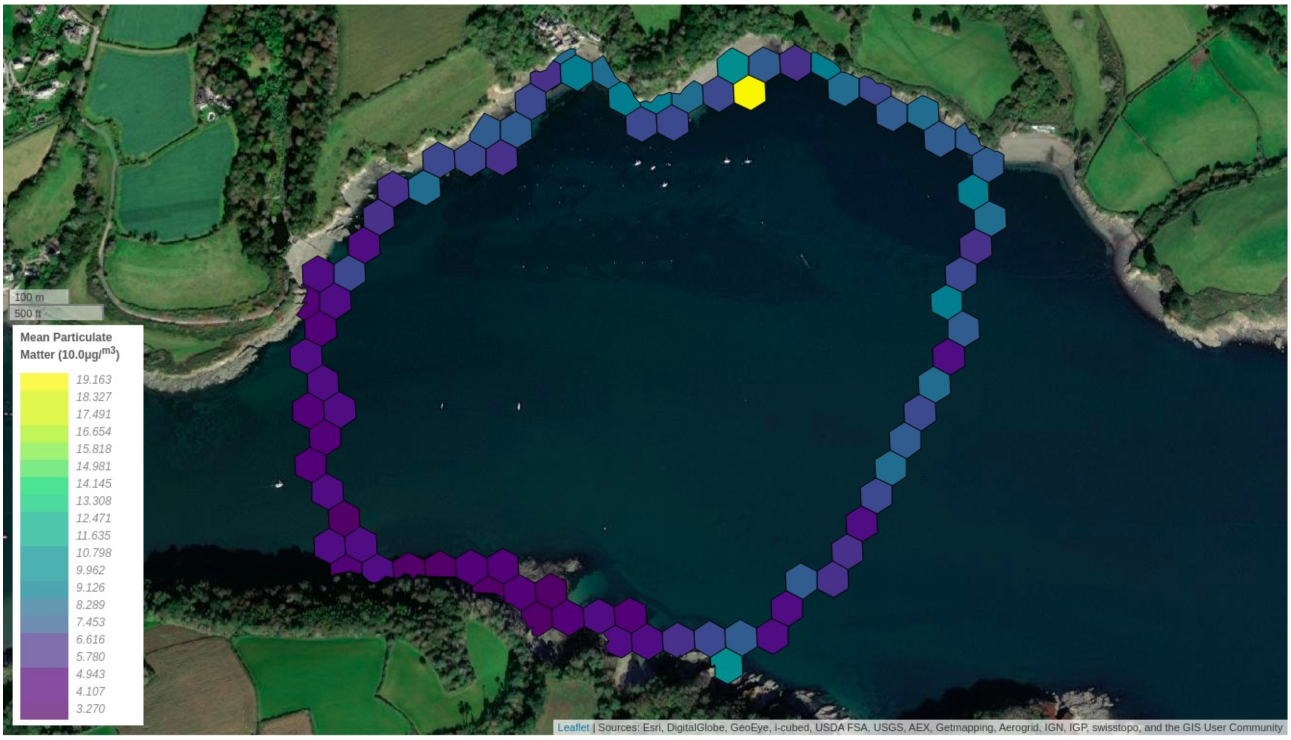


Fig. S5: Helford River, particulate matter $PM_{10}$

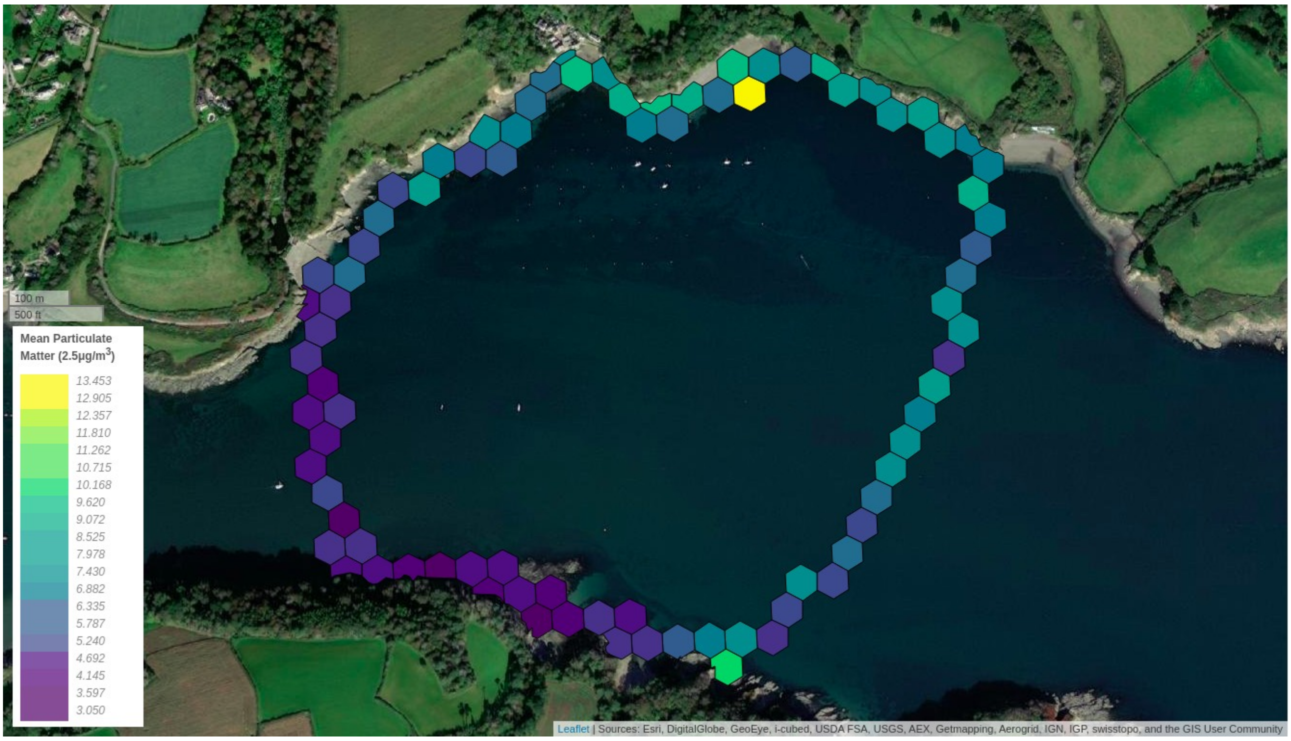


Fig. S6: Helford River, particulate matter $PM_{2.5}$

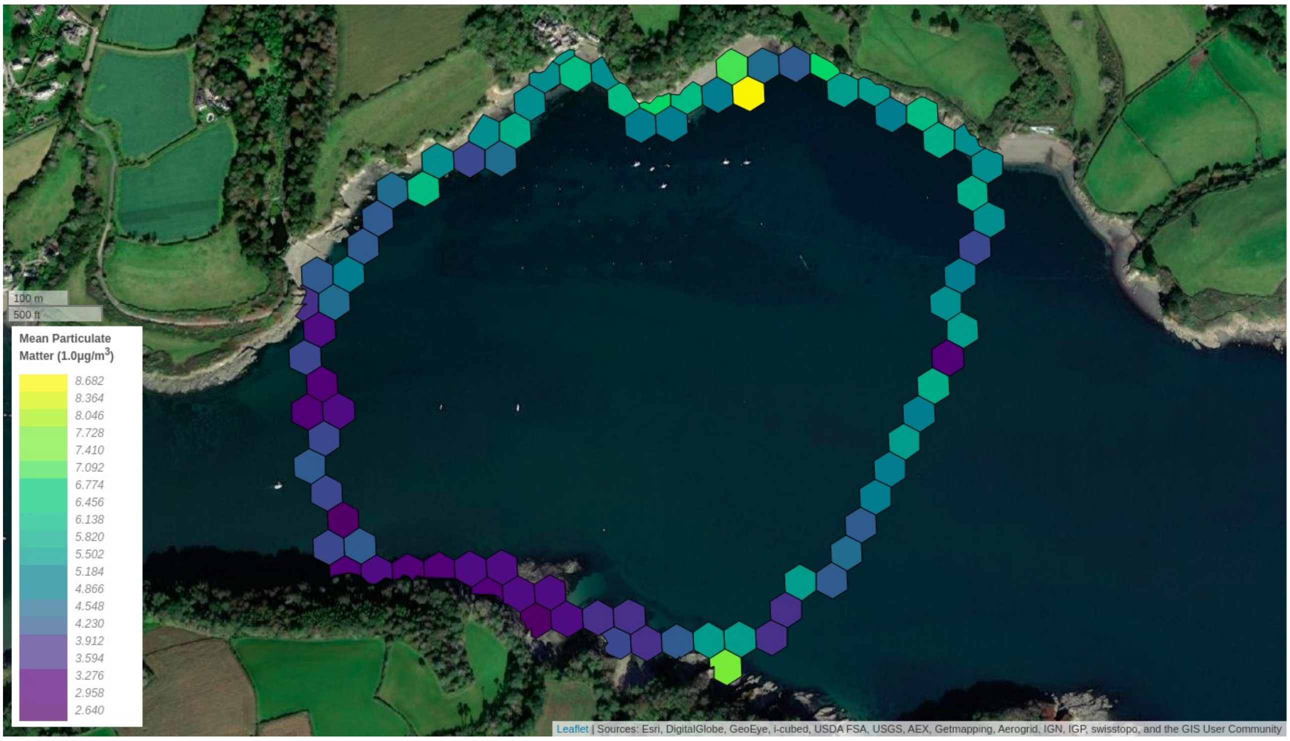


Fig. S7: Helford River, particulate matter $PM_1$

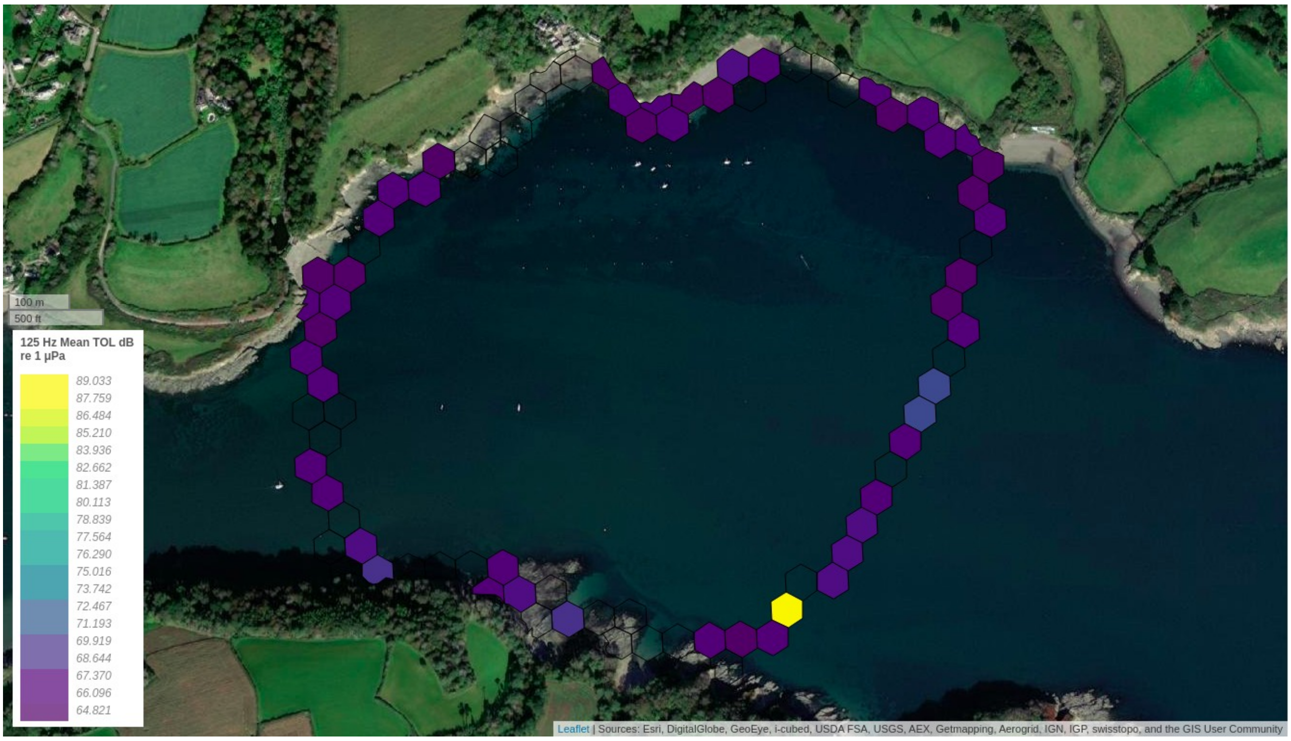


Fig. S8. Helford River, 125 Hz sound.

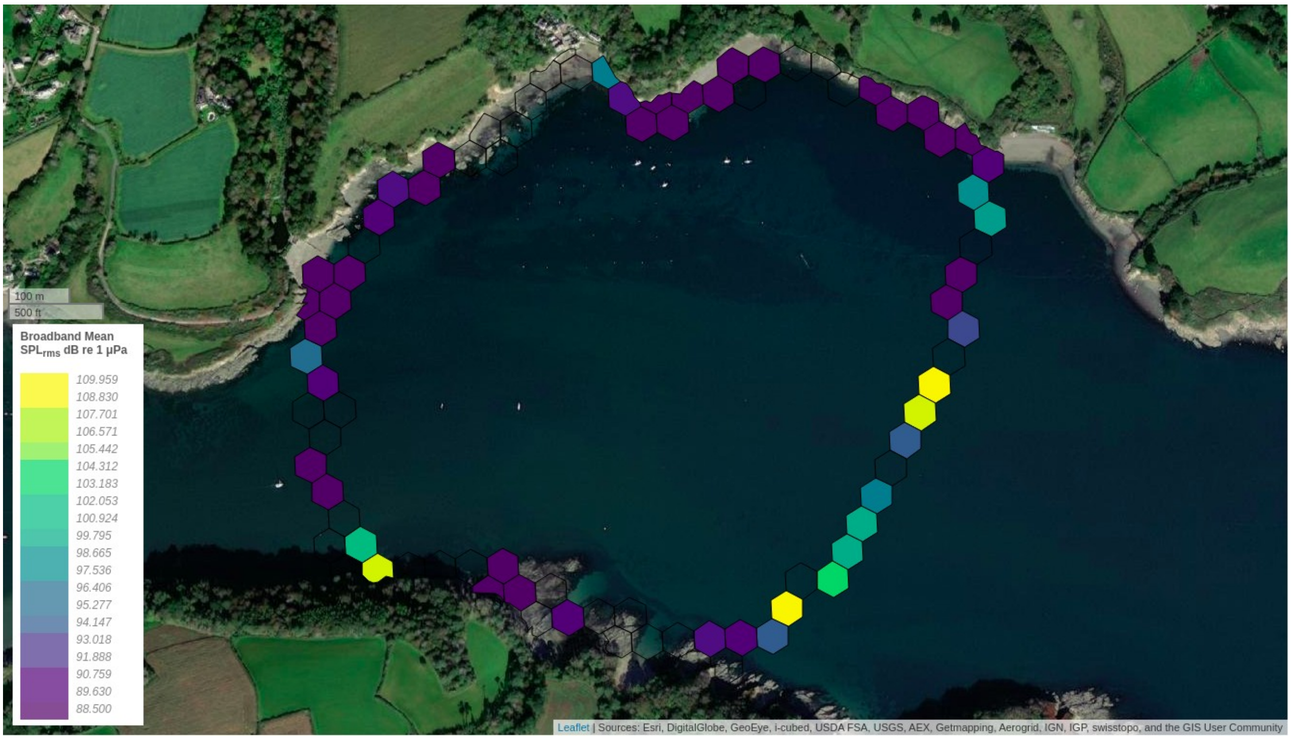


Fig. S9: Helford River, broadband mean $SPL_{rms}$

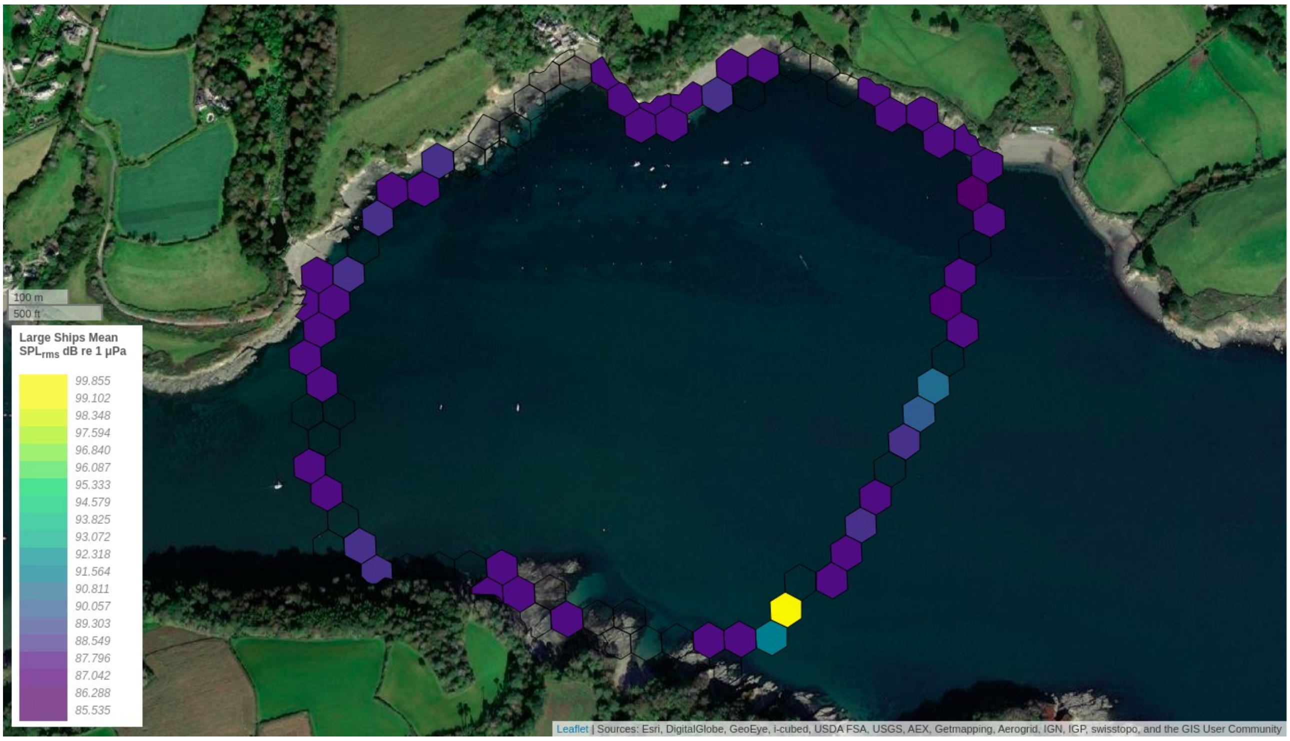

Fig. S10: Helford River, sound from large ships

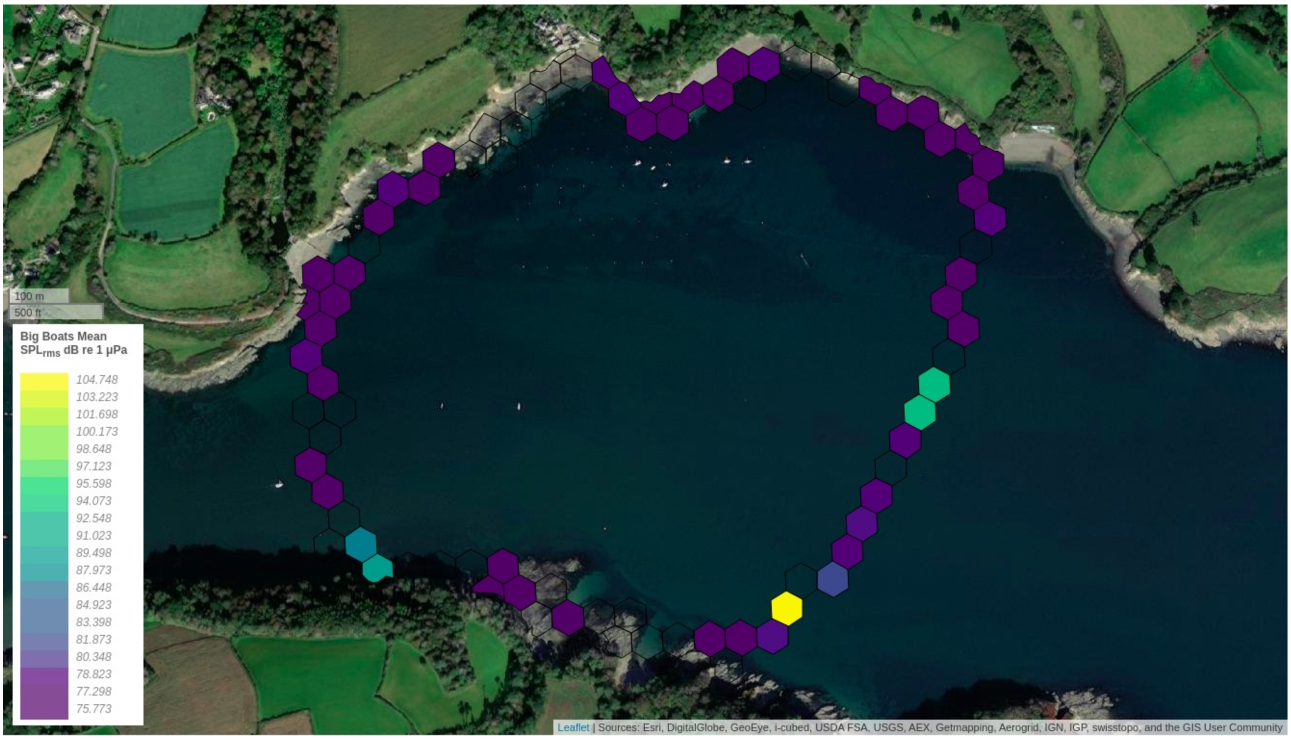

Fig. S11: Helford River, sound from big boats.
